# Concussion decreases brain signal variability

**DOI:** 10.64898/2026.11.05.764609

**Authors:** Douglas H. Schultz, Heather C. Bouchard, Bethany C. Barnwell, Nathan M. Muncy, Ramsey R. Wilcox, Cary R. Savage, Maital Neta, Aron K. Barbey

**Author notes:** Corresponding authors: Douglas H. Schultz, University of Nebraska-Lincoln, C79 East Stadium, Lincoln, NE 68588; Aron K. Barbey, University of Notre Dame, E312 Corbett Family Hall, Notre Dame, IN 46556.

## Abstract

Sports-related concussion represents a major public health concern, with millions of cases diagnosed annually. Despite significant methodological advances in functional brain imaging, a clinically meaningful neuroimaging biomarker of concussion has yet to be established. This may be due in part to a reliance on traditional measures. For example, approaches examining mean activation across trials reduce responses to a single measure of central tendency and therefore fail to capture important individual differences. In contrast, blood oxygenation level dependent (BOLD) variability offers a promising new approach to measuring brain function, Specifically, it captures individual differences in the temporal variability in the BOLD signal over time. Further, it is sensitive to neural, vascular, and neurovascular coupling changes that accompany concussion. We collected baseline (prior to injury) and post-concussion (within 48 hours of injury) neuroimaging data in 56 collegiate athletes. Within this longitudinal dataset, we investigated changes in BOLD variability across multiple spatial scales. In addition, we examined their relationships with self-reported symptoms and cognitive performance. As expected, symptoms increased following concussion while computerized cognitive performance declined. BOLD variability decreased from baseline to post-concussion, particularly in the somatomotor and language networks. Moreover, changes in BOLD variability were related to changes in symptom reporting and cognitive performance. Together, these findings advance our understanding of how concussion affects brain function and highlight BOLD signal variability as a promising biomarker to aid ongoing efforts to improve the diagnosis and treatment of sports-related concussion.

## Introduction

Each year, approximately 3.8 million concussions are diagnosed in the United States (Langlois et al., 2006). Sports-related concussions account for around 300,000 of these cases annually (Coronado et al., 2015), with high-impact sports like football and soccer significantly contributing to the incidence of such injuries (Pierpoint C Collins, 2021). Diagnosing a concussion often involves a multifaceted evaluation, where clinicians gather a detailed history, conduct a neurological exam, assess self-reported symptoms, and evaluate cognitive performance (Patricios et al., 2023). While these assessments are valuable for diagnosis, they often lack the sensitivity to detect subtle changes in brain pathophysiology both at the time of injury and throughout the recovery process (McCrea et al., 2015; Tabor et al., 2023).

During the hypothesized metabolic cascade following a concussion, ion flux and the release of neurotransmitters can lead to cellular energy deficiencies. This results in metabolic uncoupling or a mismatch between energy supply and demand (Giza C Hovda, 2014). Additionally, biomechanical forces may compromise structural integrity, contributing to disruptions in brain function. Neuroimaging techniques, such as functional magnetic resonance imaging (fMRI), offer a promising avenue for identifying potential biomarkers that may be more sensitive to these functional and structural consequences of concussion than are current clinical assessments.

Historically, functional neuroimaging studies of concussion have focused on changes in functional activity observed at the group level. These approaches have examined central tendency measures of fMRI task activation and functional connectivity (Cook et al., 2020; Dogra et al., 2024; Morelli et al., 2021). While these approaches have been informative, inconsistencies in the literature have limited the utility of neuroimaging data in both a clinical context and in understanding the pathophysiology of the injury. Some of these inconsistencies may be due to the complexities of the blood oxygen level dependent (BOLD) signal used in functional neuroimaging. For example, the BOLD signal captures a myriad of changes in the brain, including changes in neural activity as measured by local field potentials (Logothetis, 2003; Logothetis et al., 2001; Magri et al., 2012), and changes in cerebral blood flow (Ances et al., 2008; Buxton, 2010). Changes in neurovascular coupling following moderate or severe traumatic brain injury have been reported in laboratory animal models (Jang et al., 2017; Toth et al., 2016), and although less is known about mild traumatic brain injuries in humans, it has been hypothesized that similar disruptions may occur (Tan et al., 2014).

Accumulating evidence suggests that BOLD signal variability is a promising neuroimaging measure for the detection of pathophysiology in multiple clinical contexts. For example, changes in BOLD variability have been observed in Alzheimer’s disease (Scarapicchia et al., 2018; Tuovinen et al., 2020), stroke (Kielar et al., 2016), post-traumatic stress disorder (Ke et al., 2015), attention-deficit/hyperactivity disorder (Nomi et al., 2018), and autism spectrum disorder (Easson C McIntosh, 2019; Sigar et al., 2025). While BOLD signal variability may provide evidence for a wide range of clinical outcomes, it may be particularly relevant to the context of concussion. Concussion has been associated with disruptions in neural activity (Brooks et al., 2018; Conley et al., 2019; Wolf et al., 2017), vasculature (Chodobski et al., 2011; Lu et al., 2021), and neurovascular coupling (Churchill et al., 2019; Jang et al., 2017; Purkayastha et al., 2019). BOLD signal variability has been associated with several physiological changes in hemodynamic function and neurovascular coupling related to changes in cerebral perfusion (Amen et al., 2016; Soustiel C Sviri, 2007), reactivity of microvessel walls (Ueda et al., 2006), and the size and density of capillaries (Park et al., 2009). BOLD signal variability is sensitive to similar physiological changes that occur following concussion which makes it particularly valuable to understand concussion.

Additional research on BOLD variability has examined developmental changes, and its relationship with cognition. These studies have largely found decreases in BOLD variability in older adults (Garrett et al., 2010; Kielar et al., 2016; Nomi et al., 2017). In addition, these decreases in BOLD variability have generally been related to worse cognitive performance (Garrett et al., 2011; Grady C Garrett, 2014), albeit in complex ways (Armbruster-Genç et al., 2016; Goodman et al., 2024). As such, the precise functional consequence of decreased BOLD variability in the brain has not been fully elucidated. However, reduced BOLD variability has been associated with decreased oxygen and glucose metabolism, decreased cerebral blood flow, increased metabotropic receptor density, and increased excitation/inhibition ratio (Baracchini et al., 2023), lending support to its use in the investigation of concussion.

Despite an incomplete understanding of the functional consequence of decreased BOLD variability, several hypotheses have been generated. One possibility is that BOLD variability is a measure of the dynamic range of the brain to enter or maintain different neural states. Decreased variability may reflect a decreased ability to rapidly and efficiently transition between states depending on current demands (Grady C Garrett, 2014). Consistent with this hypothesis, BOLD variability is positively correlated with multiple measures of signal complexity, which can reflect information processing capacity (Wehrheim et al., 2024). Another hypothesis suggests that BOLD variability may index noise and that noise can serve to improve information processing in non-linear systems (McDonnell C Ward, 2011). Importantly, both hypotheses suggest that BOLD variability is critical for efficient information processing and cognition, where decreases in BOLD variability would result in functional deficits.

Building on this work, our prior research has examined BOLD variability in the context of concussion, observing decreases in BOLD signal variability within visual and motor cortex (relative to individuals who did not sustain a concussion; Anderson et al., 2023). These brain regions are particularly vulnerable to rotational forces, which are a common mechanism of injury in sport-related concussion (Brennan et al., 2017). The combination of BOLD variability and cognitive assessment improved concussion classification relative to cognitive performance alone. It also enhanced the ability to predict increases in self-reported symptoms (Anderson et al., 2023). These findings suggest that this method may capture unique variance not currently identified by commonly used assessment measures, potentially enhancing clinical decision-making.

To date, most neuroimaging research has employed a cross-sectional design to compare athletes who sustained a concussion with an athlete control sample, to identify physiological changes in the brain (Anderson et al., 2023; Kamins et al., 2017; McCrea et al., 2017). Several recent studies have suggested that there are considerable individual differences in functional neuroimaging data of health participants that can be related to demographics, cognitive function, or other stable traits (Hearne et al., 2016; Shen et al., 2017; Smith, 2016). Thus, cross-sectional studies make it difficult to distinguish concussion-related brain changes from potential interindividual variability across groups. A longitudinal design can account for this variability. This approach maps the time course of pathophysiological changes related to concussion, while accounting for interindividual variability. Notably, this longitudinal approach has identified global hyperconnectivity during the acute and subacute phases of injury (Bouchard et al., 2023; Kaushal et al., 2019), which may be related to reduced regional BOLD variability. While preseason baseline cognitive testing has traditionally been a recommended fundamental component of clinical concussion management practices (Broglio et al., 2014), access to baseline levels of brain function through neuroimaging data is rare. Such data provide valuable context for interpreting neuroimaging findings and understanding the underlying pathophysiology following a sports-related concussion.

The goal of the current project was to determine if concussion results in decreased BOLD variability, and if these effects underlie changes in cognitive performance and self-reported symptoms post-concussion. Importantly, our longitudinal study was designed to examine changes in BOLD variability from before to after concussion within the same participants. We hypothesized that BOLD variability would decrease from baseline to post-concussion. More specifically, we hypothesized that BOLD variability would decrease in somatomotor and visual networks, consistent with previous cross-sectional data (Anderson et al., 2023). This hypothesis was also based on the general trend of reductions in BOLD variability being associated with decreased cognitive performance (Garrett et al., 2011; Grady C Garrett, 2014).

## Methods

### Participants

We recruited 411 athletes from the University of Nebraska-Lincoln (UNL) men’s football and women’s soccer teams from 2018 until 2023. Sixty-nine of those participants were diagnosed with a concussion during the study and thus were eligible for inclusion in this longitudinal analysis. Nine participants were missing fMRI data from at least one timepoint (baseline or post-concussion), and four additional participants were excluded because they failed multiple validity indicators (Schatz C Glatts, 2013) related to differences in task performance (Higgins et al., 2024). We wanted to increase the probability that behavioral differences were explained by concussion rather than by differences in data validity. Thus, our sample included 56 athletes (men’s football n = 48; women’s soccer n = 8).

At baseline, participants ranged in age from 18 to 23 years old (mean age = 19.41; SD = 1.55). The UNL Institutional Review Board approved the project. Participants provided independent written consent for the MRI portion of the study and for using standard of care clinical data for research. Participants could consent to using both MRI and clinical data, using one but not the other, or using neither for research. All participants provided consent for using MRI data. The first year of the study (football, 2018; soccer, 2019) all participants completed baseline MRI scans. Since that time, incoming participants completed baseline scans prior to participating in their sport. Clinical baseline assessment was also performed when participants entered their sport. The assessment measured cognitive performance with the Immediate Post-Concussion Assessment and Cognitive Testing (ImPACT). This includes a baseline symptom scale, and cognitive composites based on task performance. We used verbal memory, visual memory, visual-motor speed, and reaction time composite scores in the current study. Clinical baseline measures were collected by members of the Department of Athletics concussion management team using a standardized protocol. Concussion diagnosis and management was also completed by the Department of Athletics. Following concussion diagnosis, MRI scans and clinical assessments were repeated (within approximately 48 hours of the time of injury).

Analyses including behavioral data from the ImPACT were limited to participants who had consented to using their clinical data for research and had valid ImPACT data. Twenty-two participants were missing ImPACT data (n = 14) or had not consented to using their clinical data (n = 8). The final sample for examining brain-behavior relationships across baseline and post-concussion timepoints was 36 participants.

### MRI acquisition

MRI data were collected on a 3T Siemens Skyra scanner within the Center for Brain, Biology and Behavior at UNL. At baseline and post-concussion, each participant contributed T1-weighted (TR = 2530 ms, TE = 1.69, 3.55, 5.41, 7.27 ms, flip angle = 7 degrees, FOV = 256 mm/100% phase, 176 slices, slice thickness = 1mm) and multi-band echo-planar resting-state fMRI (TR = 1000 ms, TE = 29.8 ms, flip angle = 60 degrees, FOV = 210 mm/100% phase, slice thickness = 2.5 mm, 51 interleaved slices, multi-band acceleration factor = 3). Framewise Integrated Real-Time MRI Monitoring (FIRMM) was used to evaluate real-time head motion during scans to improve data quality (Dosenbach et al., 2017). Our goal was to collect 30 minutes of low-motion data (framewise displacement < 0.2 mm), and scans were extended until either the goal was reached, or the allotted scan time was over. Some participants had one continuous resting-state scan. Other participants had their resting-state data split across two or three runs (breaks were inserted to provide feedback if participants were moving). MRI acquisition parameters and procedures were identical for baseline and post-concussion scans.

### MRI preprocessing

MRI data preprocessing was conducted with FreeSurfer (Fischl, 2012), FSL (Jenkinson et al., 2012), and AFNI (Cox, 1996). The general preprocessing approach has been described previously (Wilcox C Barbey, 2023). The T1-weighted high-resolution structural scan was brain-extracted and segmented using FreeSurfer to identify gray matter, white matter, and CSF voxels. Resting-state data were preprocessed with tools from FSL and AFNI. We used conventional preprocessing steps including motion correction, distortion correction and registration to T1 space, intensity normalization, non-aggressive ICA-AROMA denoising, nuisance signal regression, temporal bandpass filtering (0.01 – 0.1 Hz), linear detrending, and nonlinear transformation to fsaverage space.

ICA-AROMA was selected as this method performs well compared to other denoising approaches (Parkes et al., 2018). The intensity normalized functional images were smoothed using a 5 mm full width half-max kernel at a 75% brightness threshold using FSL’s SUSAN (Pruim et al., 2015). Noise components were non-aggressively removed from the non-smoothed functional images using the RegFilt function in FSL. Following the removal of motion-related noise components, white matter and CSF signals were averaged across all voxels identified with the FreeSurfer segmentation of the high-resolution T1 data. Regressors were removed from the functional images with 3dtProject in AFNI while simultaneously performing bandpass filtering and linear detrending (Lindquist et al., 2019). Finally, the denoised images were nonlinearly transformed to fsaverage space using FNIRT in FSL.

### BOLD variability

We computed resting-state mean square successive difference (MSSD) as the standard deviation of successive differences in the timeseries signal (Garrett et al., 2011; Nomi et al., 2017; Von Neumann et al., 1941) extracted from each of the 360 cortical regions identified in a multimodal parcellation atlas (Glasser et al., 2016). As participants had different numbers of runs of resting-state data, we calculated MSSD for each run and calculated a mean weighted by the number of volumes in each run divided by the total number of resting-state volumes for each participant at that timepoint. We also used well-validated network partition to assign each region to a functional network (Ji et al., 2019) (Figure 1).

**Figure 1.**
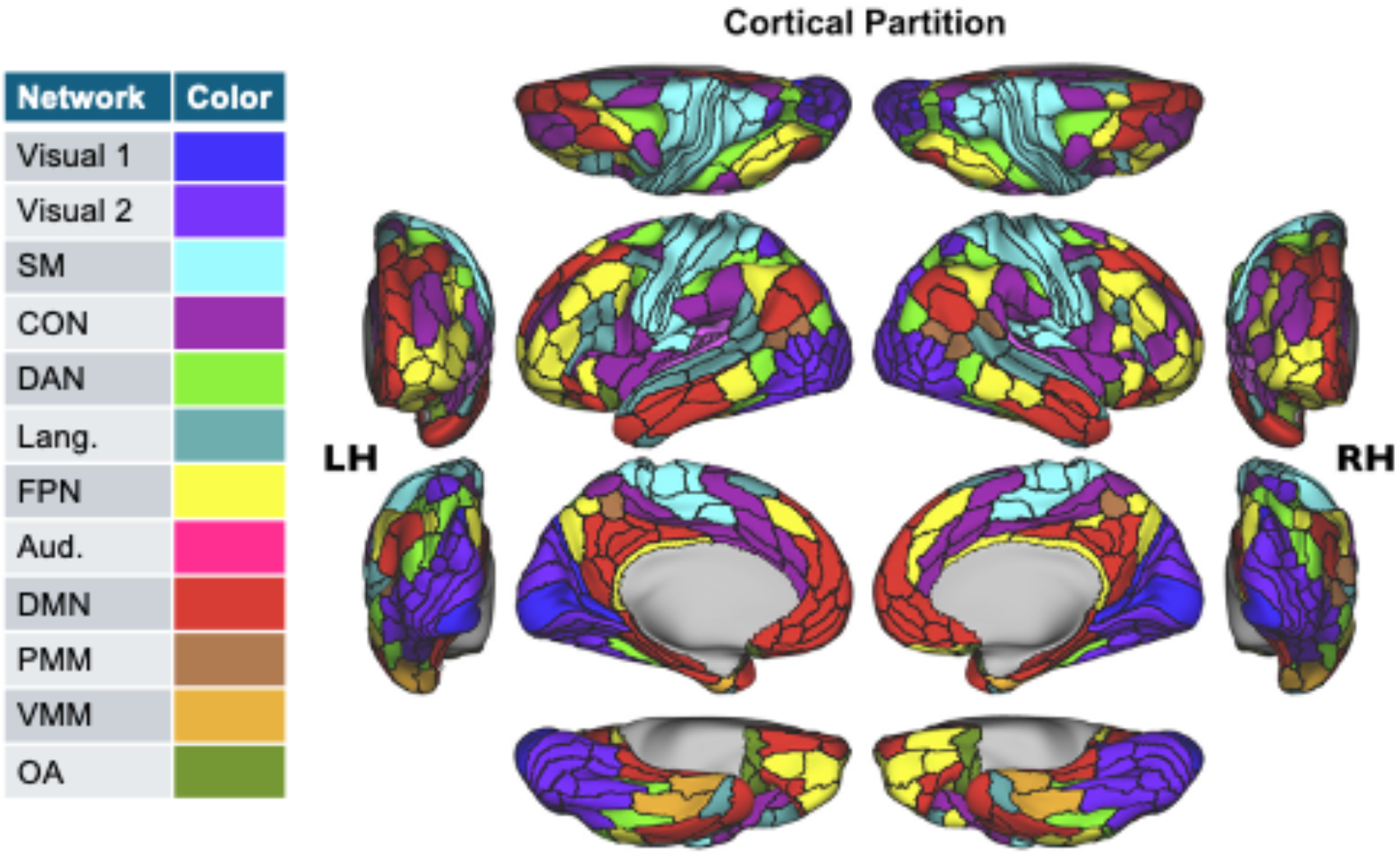
Parcellation and network assignment. SM – somatormotor, CON – cingulo-opercular, DAN – dorsal attention, Lang. – language, FPN – frontoparietal, Aud. – auditory, DMN – default mode, PMM – posterior multimodal, VMM – ventral multimodal, OA – orbito-affective.

### Data analysis

ImPACT cognitive composite scores and self-reported symptom data at baseline were compared to the same measures collected post-concussion. We examined five different symptom categories: cognitive, ocular-motor, headache-migraine, vestibular, and anxiety-mood (Lumba-Brown et al., 2020). We used the nonparametric Wilcoxon signed-rank test in JASP (Love et al., 2019) to compare baseline to post-concussion data because several of the variables were not normally distributed. We used false discovery rate (Benjamini C Hochberg, 1995) to account for multiple comparisons across the ImPACT composites and symptom measures.

We compared baseline and post-concussion BOLD variability with paired t-tests. Comparisons were made at several levels of analysis: global, network, and regional. We calculated BOLD variability on a global scale by extracting the mean timeseries signal from the whole cortex. At the network level, we calculated the mean MSSD value across regions assigned to each network for each participant and timepoint and then compared values from baseline and post-concussion with paired t-tests. We used a permutation-based approach with 1000 permutations to account for multiple comparisons across the 12 functional networks (Nichols C Holmes, 2002) as it is generally more conservative than FDR and we wanted to reduce the probability of type I error in the comparisons of BOLD variability. In networks with significant changes in BOLD variability, we compared BOLD variability on the region level. We used the same permutation-based approach to account for multiple comparisons across the total number of regions tested.

We used linear regression to examine the relationship between changes in BOLD variability and changes in the self-reported symptoms and cognitive compositive scores from the ImPACT. We used the default “Enter” entry method for linear regression in JASP. We calculated the change in BOLD variability from baseline to post-concussion for each participant in each network of interest (post-concussion – baseline). This served as the dependent variable in each of the regression models. We also calculated change scores on each of the behavioral measures of interest from the ImPACT in the same manner. These difference scores were Z-scored and served as the predictor variables in the model.

## Results

### Behavioral data

As expected, symptoms increased from baseline to post-concussion. We observed significant increases in headache-migraine (z = -3.44, FDR-corrected p = 0.005), and vestibular symptoms (z = -3.32, FDR-corrected p = 0.005). Cognitive (z = -2.32, uncorrected p = 0.02), ocular-motor (z = -1.32, uncorrected p = 0.191), and anxiety-mood (z = -0.89, uncorrected p = 0.38) symptoms all increased, but did not survive multiple comparison correction. Regarding cognitive performance, the visual memory (z = 2.2, uncorrected p = 0.029), verbal memory (z = 1.76, uncorrected p = 0.08), and reaction time (z = -1.6, uncorrected p = 0.11) composite scores decreased from baseline to post-concussion, but these differences were not statistically significant following multiple comparison correction. Finally, visual-motor speed (z = -0.06, uncorrected p = 0.96) did not change from baseline to post-concussion (Figure 2).

**Figure 2.**
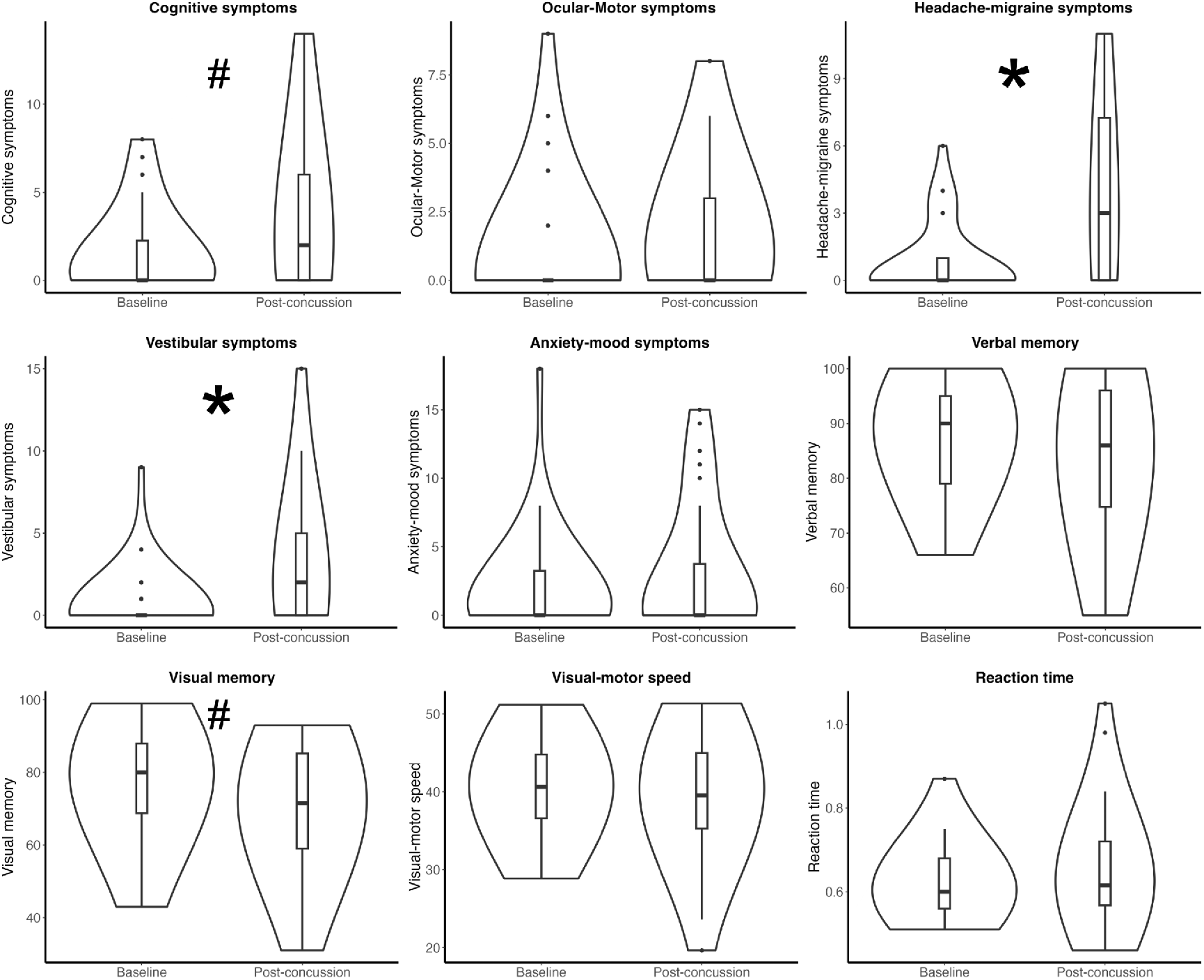
Increased symptoms and decreased cognitive performance post-concussion. Violin plots for symptoms and ImPACT cognitive composites at baseline and post-concussion. Violin plots show the distribution of scores across time points, with boxplots indicating the median and interquartile range. Points represent outliers defined as values more than 1.5 times the interquartile range from the first or third quartile. Asterisks indicate corrected p-value < 0.05. Hashmarks indicate uncorrected p-values < 0.05.

### Differences in BOLD variability from baseline to post-concussion

At a global level we observed a decrease in BOLD variability from baseline to post-concussion, *t*(59) = 2.187, p = 0.033, suggesting that BOLD signal variability across the entire brain decreased following concussion. At the network-level, we found significant decreases in BOLD variability from baseline to post-concussion in the somatomotor and language networks (Table 1). Within these networks (somatomotor and language networks), we examined BOLD variability at the region level, identifying five regions (4 in the somatomotor network and 1 in the language network) showing a significant decrease in BOLD variability from baseline to post-concussion (Figure 3).

**Figure 3.**
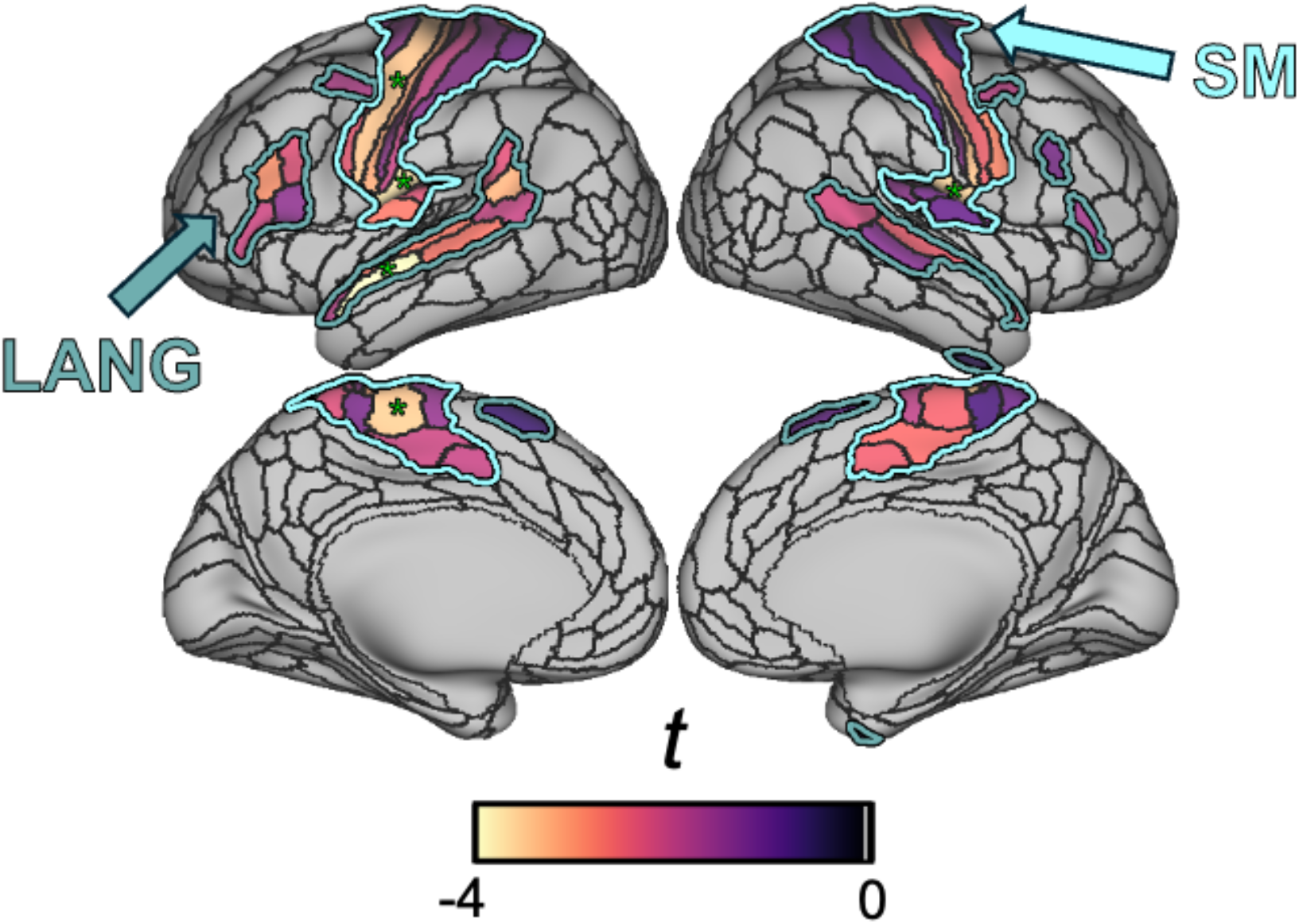
Regional decreases in BOLD variability from baseline to post-concussion. Region level comparisons of BOLD variability from baseline to post-concussion. We focused on regions in the somatomotor (regions outlined in cyan) and language (regions outlined in teal) networks as these networks demonstrated statistically significant changes. Negative values indicate a decrease in BOLD variability from baseline to post-concussion. Regions with an asterisk indicate corrected p-value < 0.05.

**Table 1.**

| **Network** | ***t*-value** |
|---|---|
| Visual 1 | -1.662 |
| Visual 2 | -1.966 |
| **Somatomotor** | **-2.814\*** |
| Cingulo-opercular | -2.290 |
| Dorsal attention | -2.107 |
| **Language** | **-3.311\*** |
| Frontoparietal | -2.212 |
| Auditory | -2.539 |
| Default mode | -1.800 |
| Posterior multimodal | -2.503 |
| Ventral multimodal | -1.341 |
| Orbito-affective | -2.177 |

Asterisks indicate corrected p-value < 0.05

### Changes in BOLD variability related to changes in behavior

We used linear regression to examine the relationship between the change in behavioral measures from baseline to post-concussion to the change in network-level BOLD variability. We used all of the behavioral measures (self-reported cognitive, oculo-motor, headache-migraine, vestibular, and anxiety-mood symptoms along with verbal memory, visual memory, visual-motor speed, and reaction time performance variables) and used the changes in those measures to predict the change in network-level BOLD variability for the somatomotor and language network. This resulted in two models, one for the somatomotor network and one for the language network. We found that the model predicting changes in BOLD variability in the somatomotor network was significant, with changes in cognitive symptoms and verbal memory as the only significant predictors (Table 2). This suggests that increased cognitive symptoms and decreased verbal memory performance from baseline to post-concussion are associated with larger decreases in BOLD variability in the somatomotor network. A similar model to predict changes in BOLD variability in the language network was not significant (Table 2).

**Table 2.**

| Model | SE | *b* | t | p |
|---|---|---|---|---|
| **Somatomotor (*F*(9,24) = 2.338, p = 0.047\*, adjusted R<sup>2</sup> = 0.267** |  |  |  |  |
| Intercept | 17.58 |  | -1.25 | 0.22 |
| **Cognitive sympt.** | **21.18** | **-0.47** | **-2.60** | **0.02\*** |
| Oculo-motor sympt. | 22.43 | 0.07 | 0.37 | 0.72 |
| Headache-migraine sympt. | 20.75 | -0.28 | -1.39 | 0.18 |
| Vestibular sympt. | 16.11 | -0.27 | -1.50 | 0.15 |
| Anxiety-mood sympt. | 15.52 | -0.2 | -1.02 | 0.32 |
| **Verbal memory** | **1.15** | **-0.36** | **-2.09** | **0.047\*** |
| Visual memory | 0.81 | -0.31 | -1.76 | 0.09 |
| Visual-motor speed | 2.42 | 0.09 | 0.42 | 0.68 |
| Reaction time | 143.29 | -0.08 | -0.40 | 0.69 |
| Language (*F*(9,24) = 1.106, p = 0.396, adjusted R<sup>2</sup> = 0.028 |  |  |  |  |
| Intercept | 17.14 |  | -1.71 | 0.10 |
| Cognitive sympt. | 20.66 | -0.17 | -0.80 | 0.43 |
| Oculo-motor sympt. | 21.88 | 0.02 | 0.08 | 0.94 |
| Headache-migraine sympt. | 20.24 | -0.31 | -1.32 | 0.20 |
| Vestibular sympt. | 15.71 | -0.18 | -0.85 | 0.40 |
| Anxiety-mood sympt. | 15.14 | -0.14 | -0.63 | 0.54 |
| Verbal memory | 1.12 | -0.25 | -1.26 | 0.22 |
| Visual memory | 0.79 | -0.40 | -1.96 | 0.06 |
| Visual-motor speed | 2.36 | 0.21 | 0.89 | 0.38 |
| Reaction time | 139.74 | -0.10 | -0.41 | 0.69 |

Asterisks indicate p-value < 0.05

### Post-concussion behavioral data does not predict post-concussion BOLD variability

We hypothesized that including baseline measures of BOLD variability and behavior would result in greater sensitivity to detecting brain-behavior relationships. We identified the change in BOLD variability from baseline to post-concussion was related to the changes in behavioral measures from baseline to post-concussion. We used a similar linear regression approach to examine the relationship between post-concussion BOLD variability and post-concussion behavioral measures. This approach does not include the baseline measures we collected. As such, we included all participants who had consented to using their clinical data and who had valid post-concussion behavioral data. We found that the model predicting post-concussion BOLD variability in the somatomotor network was not significant (Table 3). We also found that the model predicting post-concussion BOLD variability in the language network was not significant (Table 3).

**Table 3.**

| Model | SE | *b* | t | p |
|---|---|---|---|---|
| Somatomotor (*F*(9,31) = 0.819, p = 0.603, adjusted R<sup>2</sup> = -0.042 |  |  |  |  |
| Intercept | 198.50 |  | 0.42 | 0.68 |
| Cognitive sympt. | 8.48 | -0.38 | -0.84 | 0.41 |
| Oculo-motor sympt. | 11.77 | -0.02 | -0.05 | 0.96 |
| Headache-migraine sympt. | 6.58 | 0.49 | 1.52 | 0.14 |
| Vestibular sympt. | 7.33 | -0.14 | -0.36 | 0.72 |
| Anxiety-mood sympt. | 3.92 | 0.06 | 0.28 | 0.78 |
| Verbal memory | 1.38 | 0.20 | 0.79 | 0.44 |
| Visual memory | 1.17 | 0.23 | 0.97 | 0.34 |
| Visual-motor speed | 3.00 | -0.29 | -1.03 | 0.31 |
| Reaction time | 148.27 | 0.04 | 0.18 | 0.86 |
| Language (*F*(9,31) = 0.773, p = 0.642, adjusted R<sup>2</sup> = -0.054 |  |  |  |  |
| Intercept | 152.52 |  | 1.07 | 0.29 |
| Cognitive sympt. | 6.51 | -0.42 | -0.94 | 0.35 |
| Oculo-motor sympt. | 9.04 | -0.06 | -0.18 | 0.86 |
| Headache-migraine sympt. | 5.05 | 0.45 | 1.38 | 0.18 |
| Vestibular sympt. | 5.63 | 0.20 | 0.53 | 0.60 |
| Anxiety-mood sympt. | 3.01 | -0.26 | -1.14 | 0.26 |
| Verbal memory | 1.06 | 0.25 | 1.00 | 0.33 |
| Visual memory | 0.90 | 0.08 | 0.31 | 0.76 |
| Visual-motor speed | 2.31 | -0.33 | -1.15 | 0.26 |
| Reaction time | 113.92 | -0.02 | -0.07 | 0.95 |

## Discussion

We hypothesized that BOLD variability would decrease from baseline to post-concussion in the somatomotor and visual networks in our longitudinal sample. We also hypothesized that the magnitude of the decrease in BOLD variability would be related to the changes in cognition and self-reported symptoms from baseline to post-concussion. Concussion resulted in a decrease in BOLD variability from baseline to acute post-concussion, especially in the somatomotor and language networks. Participants reported increased headache-migraine, and vestibular symptoms. The changes in symptoms and cognitive performance were related to changes in BOLD variability in the somatomotor network.

We expected concussion-related decreases in BOLD variability in the somatomotor and visual networks. Consistent with our hypothesis, we found decreases in BOLD variability in the somatomotor network. A previous study also identified decreases in BOLD variability in somatomotor regions in a sample of participants who had been recently diagnosed with a concussion relative to an independent uninjured control group (Anderson et al., 2023). The somatomotor network supports motor coordination and vestibular function, while motor incoordination, dizziness, and balance problems are symptoms used to diagnose concussion (Silverberg et al., 2023). Additionally, despite clinical recovery from concussion, athletes have double the risk of suffering a musculoskeletal injury (McPherson et al., 2019), which may also point to dysfunction in the somatomotor network. Preliminary evidence supports this interpretation with decreased somatomotor function in people with a recent concussion relative to healthy controls (Lempke et al., 2023). Other research has identified concussion-related functional and structural brain changes in the somatomotor network (Chmielewski et al., 2021; Danielli et al., 2023).

We also expected BOLD variability in the visual network to decrease following concussion. Visual symptoms are common following concussion (Fraser C Mobbs, 2022; Silverberg et al., 2023), and a previous study examining BOLD variability found decreases in visual areas for participants who had been recently diagnosed with a concussion relative to an uninjured control group (Anderson et al., 2023). Additionally, several concussion studies have identified changes in visual network functional connectivity (Amir et al., 2021; Bai et al., 2023; Bouchard et al., 2023; Gilmore et al., 2016; Lu et al., 2022; Ruiz et al., 2024).

While we observed a decrease in BOLD variability in the visual network, it was not statistically significant. In contrast with his result, a previous study found decreased BOLD variability in visual areas for a group who had recently been diagnosed with a concussion relative to uninjured control participants. However, it is possible that differences in the samples could contribute to different findings in the visual network. Our sample was slightly younger and included female participants. Another possibility is that our approach is more sensitive to concussion-related brain changes as we used a longitudinal design which included baseline data for each person while Anderson et al (2023) compared a concussion group to an independent control group.

Concussion also resulted in a significant decrease in BOLD variability in the language network. The language network (Ji et al., 2019) is left lateralized and includes Broca’s and Wernicke’s areas which is critical for language function. Unlike the somatomotor network results, we did not have an *a priori* hypothesis about the effect of concussion on BOLD variability in the language network. However, a number of studies have identified language deficits following concussion (Banks et al., 2021; Ledwidge et al., 2022; Patel et al., 2023). Language disruptions following concussion may reflect disruptions to underlying cognitive processes that contribute indirectly to language processing. Disruptions in attention, working memory, and cognitive control are often observed following concussion, and language is also dependent on these cognitive processes. There have been few fMRI studies of concussion that have focused on the language network. This may be due to the absence of this network from many popular network neuroscience atlases (Gordon et al., 2016; Power et al., 2011; Yeo et al., 2011). Future studies should examine the language network as concussion is known to disrupt multiple cognitive functions that converge on language processing.

Changes in self-reported symptoms and cognitive performance were related to the change in somatomotor network BOLD variability from baseline to post-concussion. Our longitudinal design allowed us to identify these brain-behavior relationships which were not detected when employing a cross-sectional analysis. More specifically, we found changes in cognitive symptoms and verbal memory were significant predictors for changes in somatomotor network BOLD variability. The somatomotor network is involved in movement, sensory perception, and motor control. These functions are not frequently discussed in the context of cognition, but there are several factors that may explain the observed relationship between the concussion-related change in BOLD variability in the somatomotor network and cognitive symptoms. Less variability in the BOLD signal may suggest reduced neural flexibility and adaptability making it more difficult to quickly process information and respond to stimuli. Consistent with this, “feeling slowed down” is one of the specific symptoms included in the cognitive symptom cluster. Reduced variability in the BOLD signal may also reflect impaired communication between regions involved in motor control and sensory integration. This could lead to difficulties in coordinating movements and maintaining balance. Further, impaired sensory processing, a major function of the somatomotor network, can impact complex tasks. For example, if sensory information is not processed efficiently, it can lead to difficulties in understanding and responding to complex stimuli. Consistent with this, “difficulty concentrating” is one of the specific symptoms included in the cognitive symptom cluster. Finally, functional connectivity in the somatomotor network can be used to predict cognition (Chen et al., 2022), fluid intelligence (Greene et al., 2018), and transdiagnostic psychopathology (Kebets et al., 2019), which further supports the link between somatomotor network function and complex cognition.

Our longitudinal design allowed us to detect a relationship between changes in behavior and changes in BOLD variability from baseline to post-concussion. When a cross-sectional approach was used to examine the relationship between brain and behavior only at the post-concussion time point, brain-behavior relationships were not significant. This increase in sensitivity afforded by a longitudinal design is consistent with recent findings examining functional connectivity in the context of concussion by comparing baseline data to post-injury data (Bouchard et al., 2023; Kaushal et al., 2019). While baseline neuropsychological data is frequently collected, access to baseline neuroimaging data in participants who go on to have a concussion is rarer. Future studies may want to consider the benefits of collecting baseline neuroimaging data, as it appears to improve the strength of brain-behavior relationships.

### Limitations

The current study has some limitations. First, most of the sample was male football players. Future studies should increase the sampling of female participants as they are under-represented in studies used to guide clinical care (D’Lauro et al., 2022). Increasing the sampling of female participants will improve the probability that results are generalizable and provide additional clarity on potential sex differences. Second, the sample size of the study was 56, which is relatively small. We believe that the inclusion of baseline behavioral and neuroimaging data increased our ability to detect concussion-related changes, but there are many advantages to larger samples. Prior work has suggested that larger samples increase reliability of task-based fMRI data (Turner et al., 2018) and relationships between behavior and brain features (Marek et al., 2022). However, others have suggested that different approaches can improve reliability and sensitivity with smaller samples (Lee et al., 2025; Spisak et al., 2023). It is unclear how much these factors would influence BOLD variability. The increase in consortium-level data collection efforts has clear benefits for advancing brain research and these efforts should be translated to the study of concussion. Third, BOLD variability can be influenced by several factors. Changes in neural activity, vasculature, and neurovascular coupling can all impact the BOLD signal. Future studies may want to specifically examine whether changes to one or more of these factors underlie the observed change in BOLD variability. We believe the sensitivity of BOLD variability to many overlapping physiological changes following concussion makes it a particularly useful candidate for studying the neural effects of concussion.

## Conclusion

Our findings suggest that concussion results in decreased BOLD signal variability in the somatomotor and language networks. In addition, this change in BOLD variability was associated with changes in symptoms and cognitive performance. Participants reported significantly more cognitive, vestibular, and headache symptoms following concussion along with a significant decrease in visual memory performance. Global BOLD variability decreased from baseline to post-concussion with the somatomotor and language networks exhibiting significant decreases on the network-level. Changes in symptoms and cognitive performance were significantly associated with changes in BOLD variability in the somatomotor network. Notably, these effects were evident only in the context of this longitudinal approach that includes a baseline (pre-concussion) scan within participants; no significant associations were observed in the cross-sectional, post-concussion data.

Our findings partially replicate previous work examining BOLD variability in a sample of recently concussed participants relative to an uninjured control group. We extend this work by using a longitudinal design with greater sensitivity to concussion-related changes in BOLD variability within the same participants. Our results suggest that BOLD variability may be a candidate for developing a neuroimaging biomarker for concussion and emphasize the benefits of longitudinal designs in concussion research.

## Acknowledgements

This collaborative project was made possible by the contributions of the staff and facilities of the Center for Brain, Biology and Behavior. We acknowledge the cooperation, collaboration, and contributions of UNL Athletics, which were integral to the project. Additionally, we would like to acknowledge the contributions of Joanne Murray to the collection of MRI data. This work was completed utilizing the Holland Computing Center of the University of Nebraska which receives support from the UNL Office of Research and Innovation, and the Nebraska Research Initiative. Funding for this study was provided by the Great Plains IDeA-CTR (1U54GM115458, awarded to MN) and the UNL Office of Research and Innovation (awarded to CS). The funders had no role in study design, data collection and analysis, decision to publish, or preparation of the manuscript.

## Notes

### Competing Interest Statement

The authors have declared no competing interest.

